# Targeting Human Retinoblastoma Binding Protein 4 (RBBP4) and 7 (RBBP7)

**DOI:** 10.1101/303537

**Authors:** Megha Abbey, Viacheslav Trush, Elisa Gibson, Masoud Vedadi

**Author notes:** To whom correspondence should be addressed, Masoud Vedadi.

## Abstract

RBBP4 and RBBP7 (RBBP4/7) are highly homologous nuclear WD40 motif containing proteins widely implicated in various cancers and are valuable drug targets. They interact with multiple proteins within diverse complexes such as NuRD and PRC2, as well as histone H3 and H4 through two distinct binding sites. FOG-1, PHF6 and histone H3 bind to the top of the donut shape seven-bladed β-propeller fold, while SUZ12, MTA1 and histone H4 bind to a pocket on the side of the WD40 repeats. Here, we briefly review these six interactions and present binding assays optimized for medium to high throughput screening. These assays enable screening of RBBP4/7 toward the discovery of novel cancer therapeutics.

## Introduction

Retinoblastoma binding protein 4 (RBBP4) and 7 (RBBP7) also known as Retinoblastoma Associated protein 48 (RbAp48) and 46 (RbAp46), respectively, are highly homologous nuclear WD40 motif containing proteins^1, 2^ RBBP4 and 7 (RBBP4/7) have been widely implicated in various cancers and are valuable drug targets. Elevated levels of RBBP7 in nonsmall cell lung cancer, and inhibition of migration ability of lung cancer cells upon RBBP7 knockdown have been reported. RBBP7 is also overexpressed in other cancers such as renal cell carcinoma, medulloblastoma and breast cancer^4–6^. Upregulation of RBBP7 in bladder cancer results in a cell invasion phenotype, and a decrease in its expression inhibited cell invasion. RBBP4 is also overexpressed in several cancers including hepatocellular carcinoma^8^, and acute myeloid leukemia^9, 10^. Reduction of RBBP4 expression using siRNA resulted in the suppression of tumorigenicity. In human thyroid carcinoma, RBBP4 expression is upregulated and its downregulation suppressed tumorigenicity^11^. RBBP4 suppression in glioblastoma cells also enhanced sensitivity towards temozolomide chemotherapy^12^.

RBBP4 and RBBP7 are highly homologous proteins with similar structures (89% sequence identity). They are donut shaped proteins with a seven-bladed β-propeller fold typical for WD40-repeat proteins. Distinct features include a prominent N-terminal α-helix, a negatively charged PP-loop and a short C-terminal α-helix. Two binding sites have been identified on these two proteins, a c-site on the top of the WD40 repeats (Fig.1A)^13–15^ and another located on the side of the WD40 domain between the N-terminal α-helix and PP-loop(Ser-347 to Glu-364 in RBBP7)(Fig. 1B)^15–17^. RBBP4/7 are components of several multi-protein complexes involved in chromatin remodeling, histone post-translational modifications and regulation of gene expression suggesting diverse functions for RBBP4/7. Such complexes include nucleosome remodeling and deacetylase (NuRD) complex^18^, switch independent 3A (Sin3A)^19^, and polycomb repressive complex 2 (PRC2) ^20,21^. Some interaction partners of RBBP4/7 include Metastasis Associated Protein 1 (MTA1)^18,22^, also a component of NuRD complex, and suppressor of zeste 12 (SUZ12) ^15,21^ within the PRC2 complex. Structural studies have shown that these interactions along with binding to histone H4^17^ are through the side pocket on the WD40 domain of RBBP4/7 and Nurf55 *(Drosophila* RBBP4/7)(Fig. 1B). In addition, the C-terminal α-helix may be involved in binding of peptides to this site. This pocket is unique to RBBP4/7. It has both negatively charged and hydrophobic areas (Fig. 2). As a result, hydrophobic interactions play important roles in Su(z)12-Nurf55 binding. It has been shown that binding of H4, MTA1 and Su(z)12 to RBBP4/7 and Nurf55 are mutually exclusive^15, 16^. Interestingly, RBBP4/7 are able to bind to MTA1, H3 or FOG-1 (Friend of GATA-1) independently. However, within the NuRD complex, the presence of MTA1 can destabilize the interaction of RBBP4/7 with histones H3-H4^16^.

**Figure 1.**
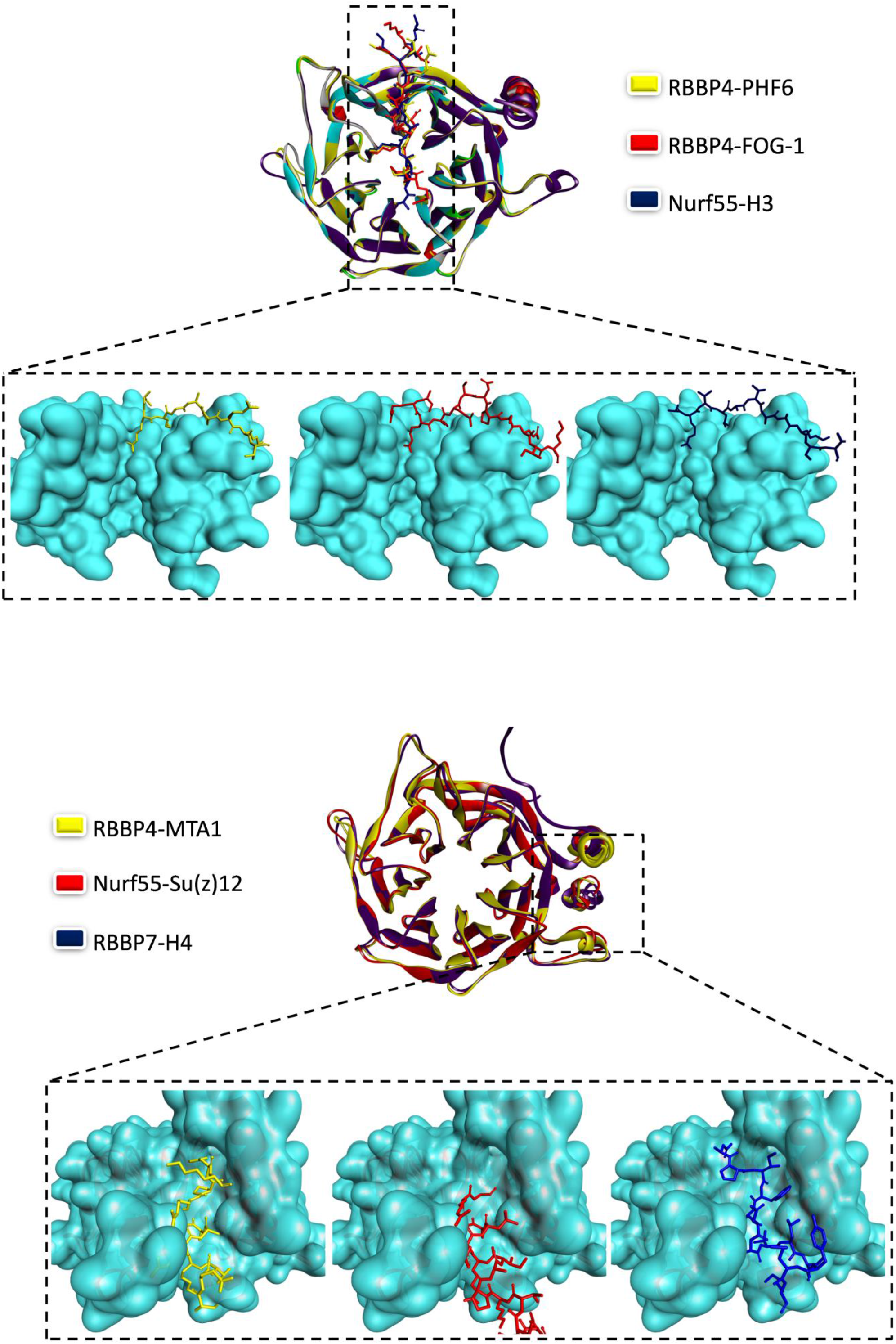
Structural details of RBBP4/7 interactions. (A) Structural alignment of RBBP4 in complex with PHF6 and FOG-1 (PDB code: 4R7A and 2XU7, respectively), and Nurf55 in complex with histone H3 peptide (PDB code: 2YBA). (B) Structural alignment of RBBP4 in complex with MTA1 (PDB code: 4PBY), RBBP7 in complex with histone H4 (PDB code: 3CFV), and Nurf55 in complex with Su(z)12 (PDB code: 2YB8). All crystal structures were downloaded from RCSB Protein Data Bank and analyzed with Accelrys Discovery studio 4.1 visualizer.

**Figure 2.**
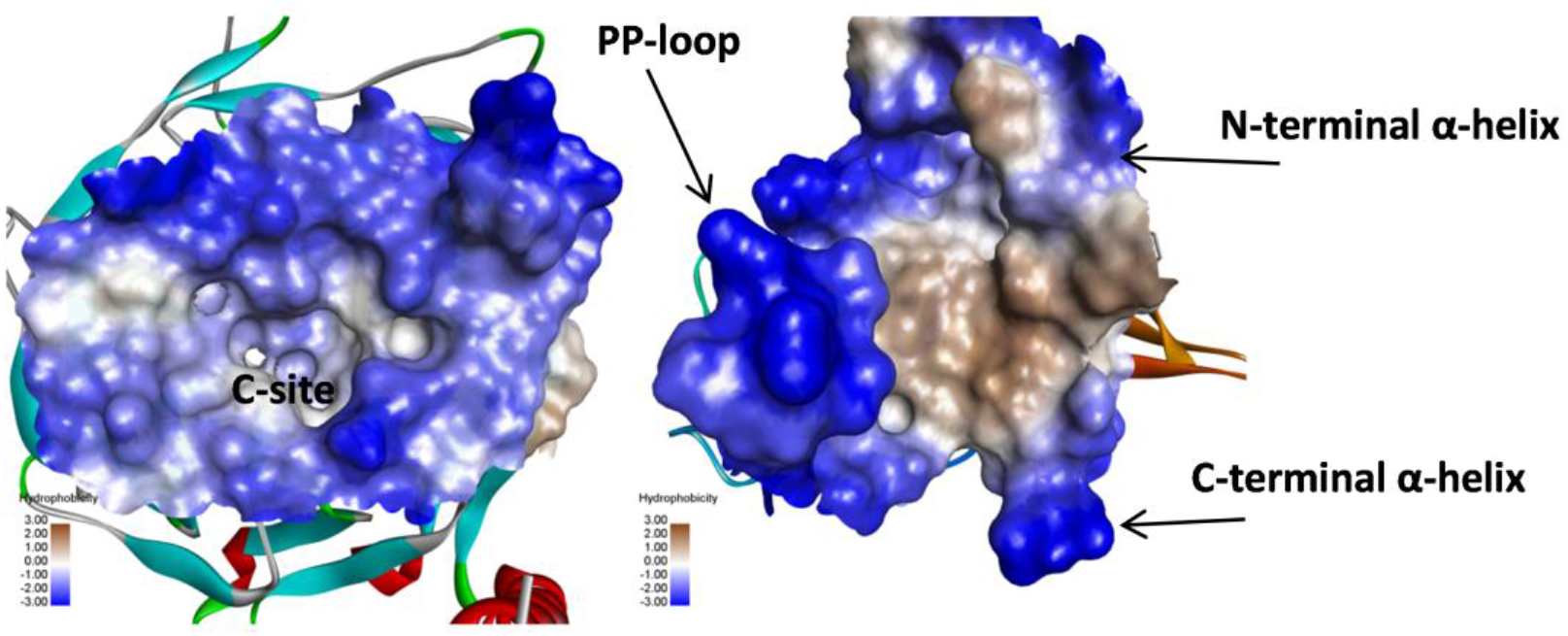
Two binding pockets on RBBP4/7. The negative charges and hydrophobic surface in the protein-binding pockets of the RBBP4/7. C-site on the *left* and side binding pocket on the *right*.

FOG-1, PHF6 (Plant homeodomain finger protein 6) and histone H3 bind to the top of the seven-bladed β-propeller fold. Crystal structures of RBBP4 in complex with PHF6^13^, FOG-1^14^, and Nurf55 with histone H3^15^ peptides reveal these ligands bind to the same position in a c-site on the top of the WD40 repeats, forming multiple hydrogen bonds. This site is negatively charged, creating a surface perfectly suited for interaction with positively charged residues of the peptides (Fig. 2). In all cases, one arginine of the peptide (Arg-2 in H3, Arg-163 in PHF6 and Arg-4 in FOG-1) is directly inserted into the negatively charged central channel of RBBP4/7 and forms a hydrogen bond with Glu-231 (Glu-235 in Nurf55) (PDB ID: 2YBA, 4R7A and 2XU7). Furthermore, all peptides adopt the same binding pose (Fig. 1A). This observation suggests that PHF6, FOG-1 and H3 interactions with RBBP4/7 and Nurf55 are highly conserved.

## Interacting proteins

### MTA1

The NuRD complex represses the transcription of several genes. It consists of several proteins, HDAC1/2 (histone deacetylase), CHD3/4/5 (chromodomain-helicase-DNA-binding protein), MBD2/3 (methyl-CpG-binding domain protein), MTA1/2/3, RBBP4/7 and Gata2a/2b (GATA binding protein)^23,24^. MTA1, 2 and 3 do not co-exist in the NuRD complex but that complex composition could rather be cell-type specific^25–28^. MTA acts as a scaffolding protein and directly interacts with HDAC1/2 and RBBP4/7. The ELM2 domain of MTA1 mediates dimerization, and its ELM2-SANT domain is responsible for recruiting HDAC1/2. The C-terminus region contains binding sites for RBBP4/7 and CHD4 (reviewed in ^29^).^18^

MTA1 has been reported to be a highly dysregulated oncogene contributing to cancer progression and metastasis (reviewed in^30^). It is upregulated in various types of cancers, including breast, ovarian and colorectal cancer, hepatocellular carcinoma and B-cell lymphomas^31^. MTA1 is a member of the MTA family and similar to MTA2 and 3, it consists of several alternatively spliced forms (reviewed in ^32^). So far, studies have largely been conducted with the full-length proteins and the roles of the spliced isoforms are yet to be explored in detail^31,33^. The N-terminus region of these proteins is well conserved, while the C-terminus differs more significantly. The conserved N-terminus consists of four domains, BAH (Bromo adjacent homology), ELM2 (EGL-27 and MTA1 homology), SANT (SWI, ADA2, N-CoR, TFIIIB-B) and the GATA-like zinc finger domains. The variable C-terminus, consists of a Src-homology 3 binding domain, an acidic region, and a bipartite nuclear localization sequence

(NLS; two in case of MTA1 and 2, one in case of MTA3) (reviewed in^32^). The expression of MTA1 in normal cells significantly varies from tissue to tissue, and is relatively higher in normal testis, brain, liver and kidney suggesting a role in these tissues (reviewed in ^31, 34^). MTA1 is mainly nuclear, but studies have reported its expression both in the nucleus and in the cytoplasm of normal and cancerous cells. Apart from being diffused in the cytoplasm, it has been shown to be associated with microtubules and with the nuclear envelope in some cases (reviewed in ^31^).^35^ MTA proteins have also been shown to interact with DNA and other co-regulators of the NuRD complex. Additionally, MTA1 serves NuRD-independent cellular functions. For example, it can associate with NURF (nucleosome remodeling factor) to activate gene expression. The methylation status of MTA1 has been reported to be a key factor for determining complex association, as methylated MTA1 recruited NuRD, while demethylated MTA1 recruited NURF (reviewed in ^31^).

It has been shown that MTA1 has two regions with similar sequences that can independently bind RBBP4, and recruits two copies of RBBP4 in the NuRD complex ^18,22^. Structural studies of complexes containing full-length RBBP4 and each of these two MTA1 regions show that the ‘side’ surface of the RBBP4 β-propeller is used for MTA1 interaction^16, 18^. Testing MTA1 peptides with various lengths (656–686, 670–695 and 670–711) for binding to full length RBBP4 by isothermal calorimetry resulted in K_D_ values of 2.3 ± 0.3, 0.05 ± 0.007 and 0.24 ± 0.16 µM, respectively^16^. Values for MTA1 (656–686) are provided in Table 1 for comparison.

## SUZ12

PRC2 (Polycomb Repressive Complex) catalyzes the trimethylation of histone H3 lysine 27 (H3K27), which results in transcriptional silencing^36, 37^ PRC2 consists of a core complex of three subunits, EZH1/2 (enhancer of zeste homologue 1/2, the catalytic histone methyltransferase subunit), EED (embryonic ectoderm development) and SUZ12 (suppressor of zeste 12). Additional proteins interact with the core complex, including RBBP4/7^38^. EZH2 activity requires interaction with both EED and SUZ12^39^. Overexpression of EZH2 has been reported in prostate and breast cancer which is associated with advanced stage and poor prognosis. Similarly, EZH2 is upregulated in various solid malignancies including renal cell carcinoma^40^, lung^41^, hepatocellular^42^ and pancreatic^43^ cancers. Knock-out studies performed with respect to the components of the PRC2 complex in mice and flies show that PRC2 and all its components are essential for viability as the knockouts exhibit embryonic lethality^44–46^. Thus, PRC2 is essential during the early stages of embryogenesis and it regulates the expression of several developmental genes ^47,48^. A study from Suz12 conditional knockout mice shows that Suz12 is important for the maintenance of hematopoietic stem cells and required for the development of the lymphoid lineage but not myeloid cells^49^. AEBP2 which serves as an associating component of PRC2 complex also interacts with RBBP4 and SUZ12, stabilizing the core PRC2 complex and increasing the methyltransferase activity of PRC2^39^. It has been shown that the three zinc fingers and RRK rich motif of AEBP2 interact with RBBP4.^50^ RBBP4 binds to AEBP2 (379–390) peptide with a dissociation constant of 7.6 ± 0.6 μM through the c-site on the top of the WD40 repeats.^50^ The crystal structures of heterotetrameric complex consisting of SUZ12, RBBP4, Jarid2, and AEBP2 fragments have recently been reported as well.^51^

BCL11A is another protein that interacts with PRC2 and NuRD and SIN3A complexes through binding to RBBP4. BCL11A competes with histone H3 for binding to the top binding site on RBBP4. However, unlike histone H3, the crystal structure of RBBP4 in complex with a BCL11A peptide (residues 2–16) shows additional novel interactions between BCL11A and the side of the propeller of RBBP4 through residues 14 to 16 of BCL11A.^52^

## PHF6

PHF6 is an X-chromosome encoded protein with nuclear and nucleolar localization, which is highly conserved among vertebrates. It consists of two atypical zinc-finger domains, called extended PHD (ePHD) domains (zinc knuckle, C2HC-type coordinating one zinc ion and an atypical PHD or ZaP)^53–55^. ePHD1 and ePHD2 share a sequence identity of 49%^56^. Although the canonical PHD domain can bind to histones, studies with ePHD2 have shown that it does not bind to any histone, but can bind to dsDNA in a non-sequence-specific manner ^54^ From murine studies, PHF6 has been shown to be highly expressed in the embryonic central nervous system including brain and anterior pituitary, and its localization shifting from nucleus and cytoplasm during development to exclusively nuclear in adult^57^. Knockdown studies indicate a role for PHF6 in cell proliferation and neuronal migration^58, 59^ Clinically, *PHF6* mutations have been implicated in two intellectual disability disorders, namely, Börjeson-Forssman-Lehmann Syndrome (BFLS) and Coffin-Siris Syndrome, in addition to T-Cell acute lymphoblastic leukemia (T-ALL), acute myeloid leukemia (AML), chronic myeloid leukemia (CML), hepatocellular carcinoma, B-cell acute lymphoblastic leukemia (B-ALL), myelodysplastic syndromes and myeloid neoplasmas.^60–70^ PHF6 is a potential oncogene which is overexpressed in various cancers such as colon, breast, stomach and glioma^71^. Its suppression in B-cell malignancies has been reported to impair tumor progression^72^. Lower expression of PHF6 has also been reported in some other cancers such as esophagus cancer^71^.

PHF6 interaction partners (NuRD complex, PAF-1 and UBF) exist both in the nucleoplasm and the nucleolus. From the NuRD complex, PHF6 was shown to co-purify with CHD3, CHD4, RBBP4, RBBP7 and HDAC1, wherein, PHF6 interaction with RBBP4 and CHD4 was restricted to the nucleoplasm, despite its presence in the nucleolus as well^53^. Among the various co-purified components of the NuRD complex, a direct interaction has been reported between RBBP4 and PHF6 and the interaction has been characterized both structurally and biophysically^13, 54^ A structure of full-length RBBP4 in complex with a PHF6 peptide (157–171 amino acids) showed that a negatively charged surface at the top of the RBBP4 β-propeller is essential for binding to a PHF6 peptide (a dissociation constant of 7.1 ± 0.4 µM; Table 1)^13^. Functionally, it was shown that the PHF6 region spanning amino acids 145 to 207 was sufficient for repressing the genes to which it was targeted, and its interaction with RBBP4 was required for the process^13, 54^

## FOG-1

FOG-1 is a 1006 amino acid protein that consists of nine zinc fingers, which are involved in mediating interactions with other proteins^73–76^. Within the NuRD complex, FOG-1 has been shown to directly interact with RBBP4. This interaction has been characterized biophysically and structurally. The N-terminus residues of FOG-1 (1–12 amino acids) have been shown to bind to RBBP4 with a K_D_ of 0.6 ± 0.13 µM^14^ (Table 1).

**Table 1.** Interaction of RBBP4/7 with reported side-surface and top-surface binding peptides.

| Protein | peptide | Assay | $K_D$ (μM) | Structure availability | Binding site | References |
| --- | --- | --- | --- | --- | --- | --- |
| RBBP4 | MTA1 (656-686) | ITC | $2.3 \pm 0.3$ | Yes | Side | Alqarni et. al., 2014<sup>16</sup> |
| Nurf55 | Su(z)12 (75-93) | ITC | $6.7 \pm 0.3$ | Yes | Side | Schmitges et.al., 2011<sup>15</sup> |
| RBBP7 | H4 (16-41) | Flu | 1.1 | Yes | Side | Murzina et.al., 2008<sup>17</sup> |
| Nurf55 | H4 (26-45) | ITC | $0.035 \pm 0.001$ | Yes | Side | Nowak et. al., 2011<sup>86</sup> |
| Nurf55 | H3 (1-15) | FP | $0.8 \pm 0.1$ | Yes | Top | Schmitges et.al., 2011<sup>15</sup> |
| Nurf55 | H3 (1-15) | ITC | $2.1 \pm 0.05$ | Yes | Top | Nowak et. al., 2011<sup>86</sup> |
| Nurf55 | H3 (1-28) | ITC | $1.6 \pm 0.1$ | Yes | Top | Nowak et. al., 2011<sup>86</sup> |
| RBBP4 | PHF6 (157-171) | ITC | $7.1 \pm 0.4$ | Yes | Top | Liu et.al., 2015<sup>13</sup> |
| RBBP4 | FOG-1 (1-12) | ITC | $0.6 \pm 0.13$ | Yes | Top | Lejon et.al., 2011<sup>14</sup> |

**Flu: fluorescence spectroscopy, FP: Fluorescence Polarization assay, ITC: Isothermal Titration Calorimetry.**

FOG-1 has been shown to be largely functional in the hematopoietic system, in addition to other tissues such as male gonads, heart, and intestine (reviewed in^77^). Accordingly, it is important for erythropoiesis and megakaryopoiesis and repressing myelo-lymphoid differentiation^78^. FOG-1 knockout mice are embryonically lethal due to severe anemia^79^. FOG-1 interaction with GATA1 has been well studied and reports also suggest its co-expression with GATA2, 3 and 4 (reviewed in^77^). Disruption in its interaction with GATA-1 results in familial dyserythropoietic anemia and thrombocytopenia^80^. FOG-1 has been shown to act both as a co-activator or a co-repressor for its target genes (summarized in^77^).^81^ FOG-1 is also capable of upregulating the expression of genes that are otherwise repressed by GATA3^82^. In addition to GATA factors, FOG-1 interacts with, and recruits co-repressor complexes, NuRD complex and C-terminal binding protein (CtBP)^83,83^.

## Binding to Histone H3 and H4

Structural data shows that the ‘side’ surface of RBBP7 (and Nurf55) β-propeller interacts with a H4 peptide and this interaction requires unfolding of helix 1 of histone H4. It has been suggested that this unfolding would be required when H3 is present alone or in a complex (for example, in the nucleosome, or with H3 and ASF-1) ^17,85,86^. On the other hand, an H3 peptide interacts with the ‘top’ surface of the Nurf55 β-propeller and thus, H3 and H4 peptides bind at different surfaces^15, 86^. Interactions between Nurf55/RBBP4/RBBP7 and H3 and H4 in the H3-H4 dimeric and tetrameric complexes have been extensively characterized ^17, 85–87^. Murzina and colleagues demonstrated that the K_D_ values for RBBP7 interaction with recombinant full-length H3-H4 complex and H4 peptide (16–41 amino acids) were 0.9 and 1.1 µM, respectively, as observed by fluorescence spectroscopy^17^. However, the N-terminal tail of H4 (1–15 amino acids) is not involved in these interactions^86^. Nurf55, on the other hand, interacts with H3 N-terminal tail peptide (1–15 amino acids) with a K_D_ of 0.8 ± 0.1 µM as measured by fluorescence polarization (FP)^15^. It has also been shown that modifications of the H3 peptide (1–15 amino acids), such as H3K4me3 and H3K9me3, reduced this interaction^86^. A similar observation was made for the interaction of RBBP4 with acetylated H4 peptide^87^. Apart from the N-terminal tails, H3 and H4 can interact with RBBP4 *via* other surfaces, though these interactions are weaker ^87^. A recent study using RBBP4 has reported that the protein interacts with a H3-H4 dimer but not the tetramer, and the K_D_ for the interaction was 0.61 ± 0.49 nM^87^. Though RBBP4 and RBBP7 interact with histone proteins directly (H3 and H4), this interaction is affected by other RBBP4/7 binding partners. Interaction with NuRD is believed to recruit this complex to nucleosomes.

## Developing a suite of assays for high throughput screening

As discussed, the dysregulation of RBBP4/7 have been widely implicated in cancers. However, no small molecule antagonist of their interactions with any of the interacting proteins has been reported to-date. This necessitates developing binding assays and optimizing them for high throughput screening toward the discovery of potent, selective and cell permeable small molecules to further investigate the roles they play in normal cells and in cancers. We aimed at the development of fluorescence polarization-based RBBP4/7 binding assays with the 6 peptides which have already been characterized, and crystallized in complex with RBBP4 or RBBP7; MTA1 (656–686; DVFYMATEETRKIRKLLSSSETKRAARRPYK)^16^, SUZ12 (105–121; LIAPIFLHRTLTYMSHR)^21^, H3 (1–21; ARTKQTARKSTGGKAPRKQLA; binding of peptides with different length was reported by Schmitges et. al.^15^), FOG-1 (1–15; MSRRKQSNPRQIKRS)^14^, PHF6 (157–171; KSKKKSRKGRPRKTN)^13^ and H4(16–41; KRHRKVLRDNIQGITKPAIRRLARRG)^17^(Fig. 1A and 1B). Available binding data for these peptides are summarized in Table 1. They bind to RBBP4/7 with a range of affinities from a K_D_ value of 35 nM for Nurf55 tight binding to H4 (26–45) peptide to low micromolar K_D_ for SUZ12 binding to histone H3. We attempted to develop binding assays for all these interactions and optimize them for screening in a 384-well format. Binding of each FITC-labeled peptide to RBBP4/7 was tested at various pH (Supplementary Figure 1A-6A) and buffers (Supplementary Figure 1B-6B). Binding of MTA1, SUZ12, histone H3 and H4 peptides generated enough signal to allow the development of assays close to physiological conditions (pH 7 to 8). The assays were performed using selected buffer and pH at various buffer concentrations (Supplementary Figure 1C-6C). Interestingly, the presence of up to 100 mM of salt (NaCl/KCl) did not have any effect on FP signal for peptides that bind to the side pocket (Supplementary Figure 1D-3D). However, for peptides that bind to the c-site on top of the WD40 fold, the presence of salt as high as 25 mM significantly decreased the binding (Supplementary Figure 4D-6D). These effects are consistent with the highly negatively charged c-site and the highly hydrophobic side pocket (Fig. 2). We further evaluated the effect of detergents (Triton X-100, Tween 20 and NP-40) as well as reducing agents (DTT and TCEP) (Supplementary Figure 1E and 1F through 6E and 6F). Optimum buffer conditions selected for performing each assay are shown in Table 2. Total fluorescence was also measured at various concentrations of FITC-labeled peptides to be able to select the lowest peptide concentration that generates enough signal (Supplementary Figure 1G to 6G). This is particularly important for tight interactions. The presence of up to 5% DMSO in assay buffers did not have a dramatic effect on FP signal and was well tolerated with all of the peptides (Supplementary Figure 1H to 6H).

**Table 2.** Optimized buffer conditions for the fluorescence polarization based assays for quantifying interaction of RBBP4/7 with their interacting peptides

| **Interaction** | **Buffer** | **Protein**<br>**(μM)** | **Peptide**<br>**(nM)** |
| --- | --- | --- | --- |
| RBBP4-MTA1 (656-686) | 50 mM Sodium phosphate, pH 7.5,<br>150 mM KCl,<br>0.01 % NP-40 | 0.15 | 10 |
| RBBP4-SUZ12 (105-121) | 50 mM Potassium phosphate, pH<br>7.0, 75 mM KCl, 0.005% NP-40 | 12 | 20 |
| RBBP4-H3 (1-21) | 10 mM Tris, pH 8.0,<br>0.005 % NP-40 | 3 | 5 |
| RBBP4-PHF6 (157-171) | 25 mM Tris, H 7.5,<br>0.02 % TX-100, 2 mM DTT | 0.30 | 5 |
| RBBP4-FOG-1 (1-15) | 50 mM Tris, pH 7.5,<br>0.005 % NP-40, 2 mM DTT | 1.20 | 5 |
| RBBP7-H4 (16-41) | 25 mM Sodium phosphate, pH 8.0,<br>75 mM NaCl,<br>0.02 % TX-100 | 0.30 | 5 |

Using the optimized assay conditions we assessed binding and determined the apparent KD values for the interaction of MTA1 (33 ± 13 nM), SUZ12 (6.4 ± 0.6 µM), PHF6 (201 ± 102 nM), FOG1 (488 ± 245 nM) and histone H3 (1.4 ± 0.2 µM) peptides to RBBP4 (Fig. 3). Binding of histone H4 peptide (16–41) to RBBP7 was also tested with a K_D_ value of 135 ± 12 nM (Fig. 3C). Typically, FP-based screening relies on displacement of peptide from the binding site on the target protein by small molecules that compete with peptide ^88,89^. To discriminate against the term IC_50_ value, which indicates the concentration of the compound that causes 50% reduction in activity of the target enzyme, we have previously introduced the term “K_disp_” which refers to the same concept but for displacement-based binding assays ^88–90^. To evaluate displacement of the peptides from RBBP4/7 binding sites, we used unlabeled version of each peptide to compete and reduce the signal (Fig. 4). Clear displacement of FITC-labelled peptides was observed for MTA1 (K_disp_ : 308 ± 15 nM), histone H4 (K_disp_ : 662 ± 8 nM), PHF6 (K_disp_ : 1.7 ± 0.2 µM), and FOG1 (K_disp_ : 6.8 ± 3.3 µM) peptides (Fig. 4A, 4C, 4D and 4E). All these peptides bind with K_D_ values below 500 nM (Fig. 3). For SUZ12 binding to RBBP4 with K_D_ values of 6.4 µM, we detected no displacement by unlabeled peptide up to 10 µM (Fig. 4B). This is probably because the range of concentrations used may not be high enough. However, we were unable to use higher concentrations of SUZ12 peptide due to low solubility. For histone H3 peptide, we only included a range of up to 1.5 µM (Fig. 4F) to test the displacement as the signal at higher concentrations of peptide was dramatically fluctuating.

**Figure 3.**
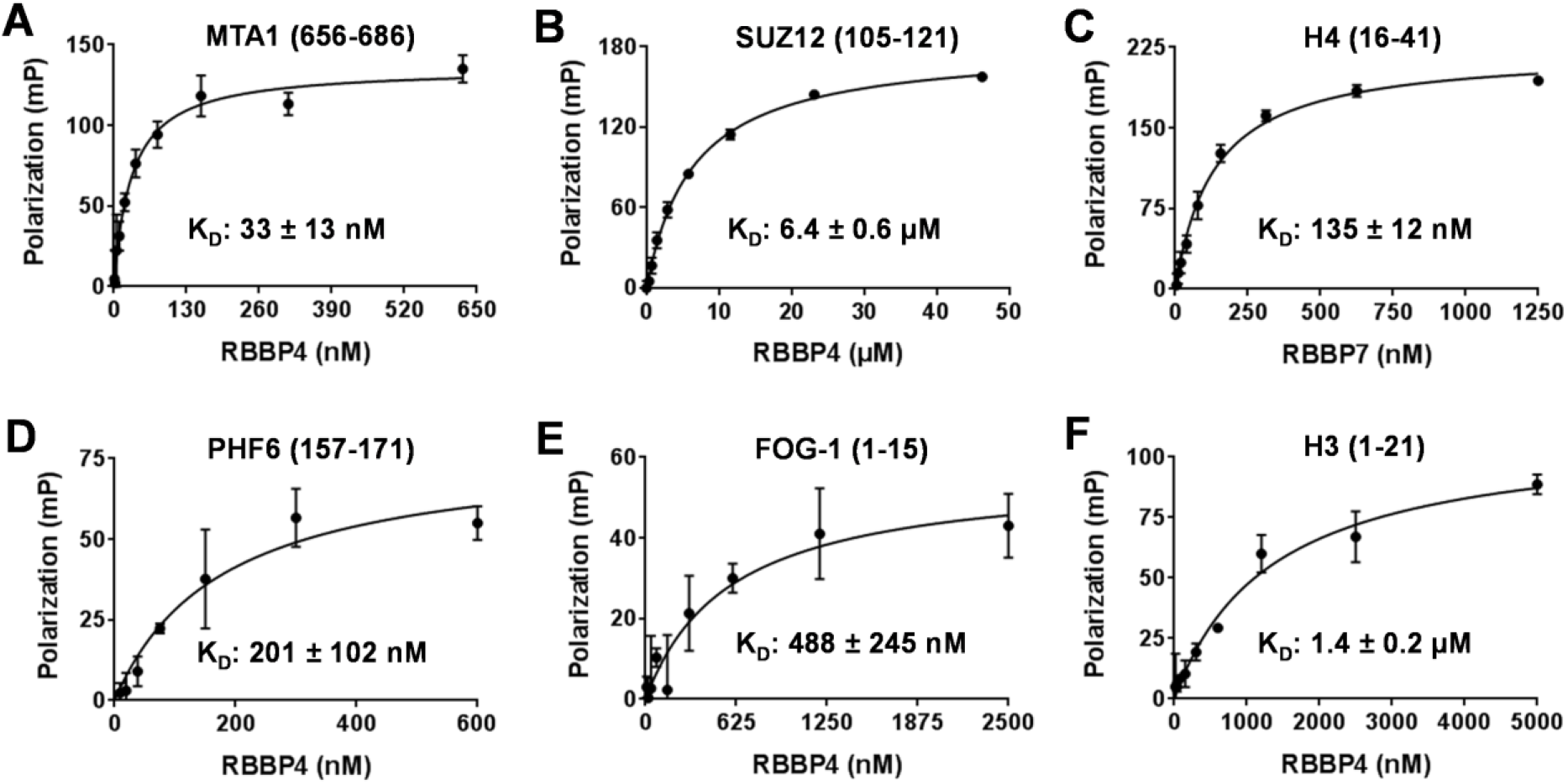
Assessing the interaction of human RBBP4/7 with peptides. Interaction of RBBP4 with FITC labelled (A) MTA1 (656–686), (B) SUZ12 (105–121), (D) PHF6 (157–171), (E) FOG-1 (1–15) and (F) histone H3 (1–21) was assessed by monitoring the increase in fluorescence polarization signal. The interaction of (C) RBBP7 with histone H4 (16–41) was assessed as well.

**Figure 4.**
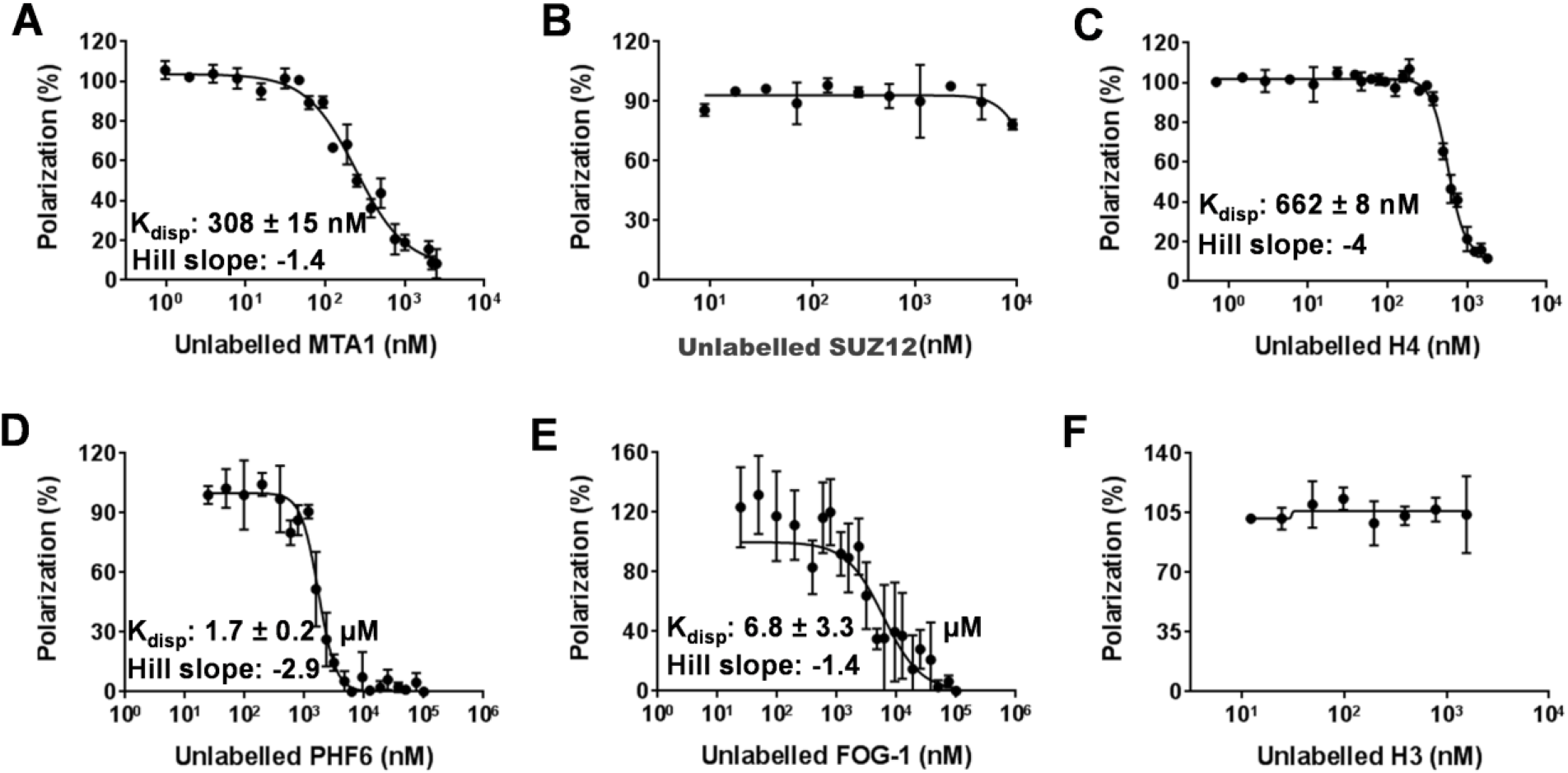
Displacement of interacting peptides. Displacement of all binding peptides shown in figure 1 using respective unlabeled peptides was assessed. FITC-labeled (A) MTA1, (D) PHF6 and (E) FOG-1 peptides were competed off RBBP4 by respective unlabeled peptides. Similarly (C) histone H4 peptide was displaced from RBBP7 binding site. Unlabeled peptide within the indicated ranges did not displace the FITC-labeled (B) SUZ12 or (F) histone H3 peptides from RBBP4 binding sites.

Although we could not show that the displacement of SUZ12 and histone H3 peptides is possible, we did not dismiss the assays for screening. Performing Z’-determination for assays with all six peptides (Fig. 5) indicated that the assay for RBBP7 with histone H4 has the highest Z’-factor (0.76; Fig. 5C) followed by RBBP4 with SUZ12 (0.63; Fig. 5B), histone H3 (0.59; Fig. 5F) and MTA1 (0.55; Fig. 5A). As expected, the assays for PHF6 and FOG-1 were not suitable for screening with Z’Factors below 0.5(Fig. 5D and 5E).

**Figure 5.**
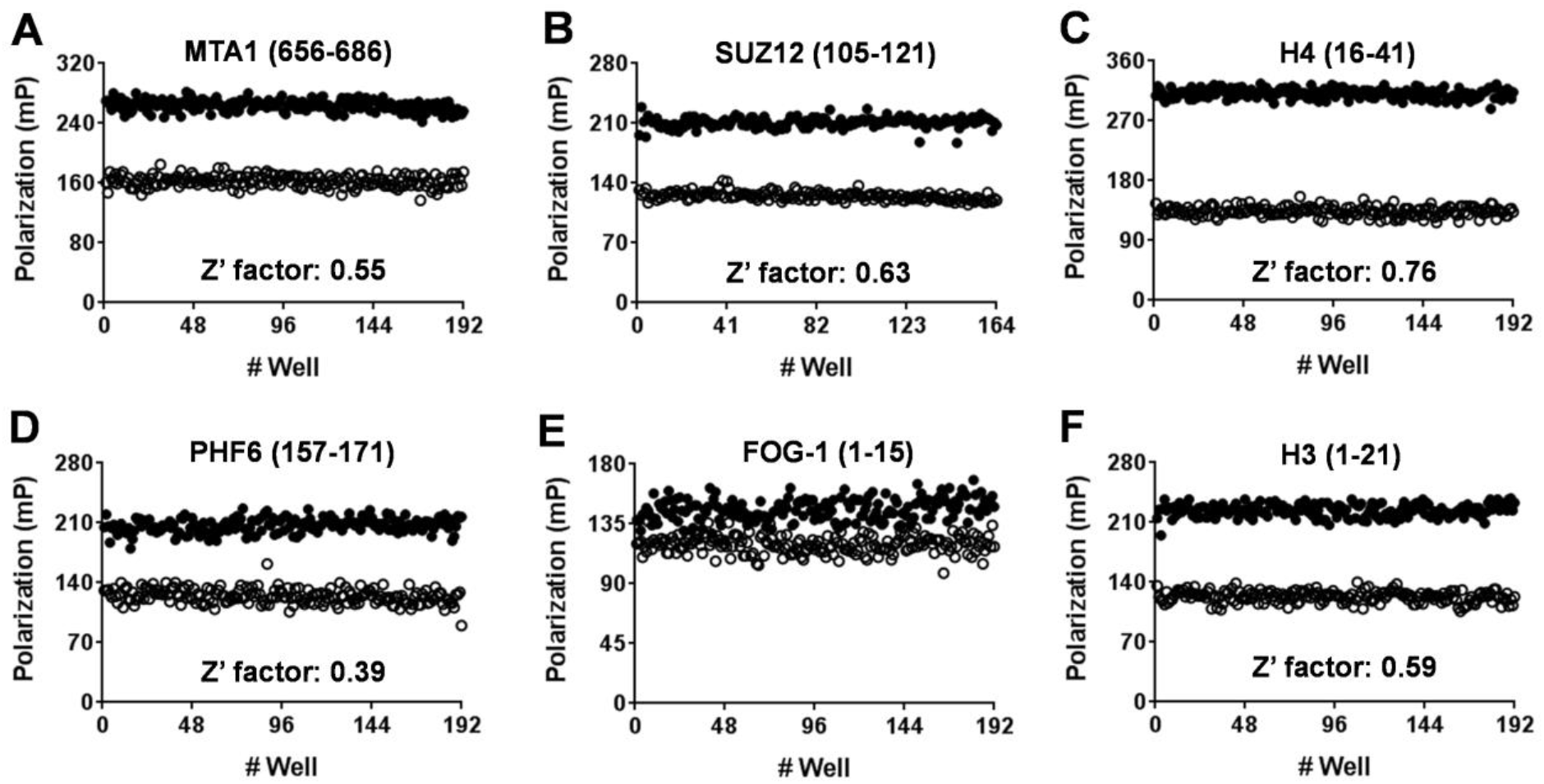
Z’-factor determination. Suitability of the peptide displacement assays for screening was determined by calculating the Z‘-factor for interaction of RBBP4 with (A) MTA1 (0.55), (B) SUZ12 (0.63), (D) PHF6 (below 0.5), (E) FOG-1 (below 0.5) and (F) H3 (1–21) (0.59). (C) Screening RBBP7 using histone H4 resulted in a Z’-factor of 0.76

In conclusion, we provide assays for displacement of MTA1, SUZ12, histone H3 and H4 peptides that have been optimized for high throughput screening of RBBP4/7, enabling discovery of small molecule antagonists that could be used toward further investigating RBBP4/7 roles in cells and discovery of cancer therapeutics.

## Contribution of authors

M.A designed and performed all experiments, analyzed data and wrote the manuscript. V.T performed structural analysis
(figure 1 and 2). E.G purified proteins. M.V led the project, designed experiments, reviewed data and wrote the manuscript.

## Acknowledgement

We would like to thank Drs. Matthieu Schapira and Peter Brown for critical review of the manuscript. The SGC is a registered charity (number 1097737) that receives funds from AbbVie, Bayer Pharma AG, BoehringerIngelheim, Canada Foundation for Innovation, Eshelman Institute for Innovation, Genome Canada, Innovative Medicines Initiative (EU/EFPIA) [ULTRA-DD grant no. 115766], Janssen, Merck & Co., Novartis Pharma AG, Ontario Ministry of Economic Development and Innovation, Pfizer, São Paulo Research Foundation-FAPESP, Takeda, and the Wellcome Trust.

## References

1. Qian, Y. W.; Wang, Y. C.; Hollingsworth, R. E. J., et al. A retinoblastoma-binding protein related to a negative regulator of Ras in yeast. Nature 1993, 364 (6438), 648–52.

2. Qian Y. W.; Lee E. Y. Dual retinoblastoma-binding proteins with properties related to a negative regulator of ras in yeast. The Journal of biological chemistry 1995, 270 (43), 25507–13.

3. Wang, C. L.; Wang, C. I.; Liao, P. C., et al. Discovery of retinoblastoma-associated binding protein 46 as a novel prognostic marker for distant metastasis in nonsmall cell lung cancer by combined analysis of cancer cell secretome and pleural effusion proteome. Journal of proteome research 2009, 8 (10), 4428–40.

4. Kim, D. S.; Choi, Y. P.; Kang, S., et al. Panel of candidate biomarkers for renal cell carcinoma. Journal of proteome research 2010, 9 (7), 3710–9.

5. Bunt, J.; Hasselt, N. A.; Zwijnenburg, D. A., et al. OTX2 sustains a bivalent-like state of OTX2-bound promoters in medulloblastoma by maintaining their H3K27me3 levels. Acta neuropathologica 2013, 125 (3), 385–94.

6. Thakur, A.; Rahman, K. W.; Wu, J., et al. Aberrant expression of X-linked genes RbAp46, Rsk4, and Cldn2 in breast cancer. Molecular cancer research: MCR 2007, 5 (2), 171–81.

7. Yeh, H. H.; Tseng, Y. F.; Hsu, Y. C., et al. Ras induces experimental lung metastasis through up-regulation of RbAp46 to suppress RECK promoter activity. BMC cancer 2015, 15, 172–185.

8. Song, H.; Xia, S. L.; Liao, C., et al. Genes encoding Pir51, Beclin 1, RbAp48 and aldolase b are up or down-regulated in human primary hepatocellular carcinoma. World J Gastroenterol 2004, 10 (4), 509–13.

9. Sakhinia, E.; Faranghpour, M.; Liu Yin, J. A., et al. Routine expression profiling of microarray gene signatures in acute leukaemia by real-time PCR of human bone marrow. British journal of haematology 2005, 130 (2), 233–48.

10. Casas, S.; Ollila, J.; Aventín, A., et al. Changes in apoptosis-related pathways in acute myelocytic leukemia. Cancer Genetics and Cytogenetics 2003, 146 (2), 89–101.

11. Pacifico, F.; Paolillo, M.; Chiappetta, G., et al. RbAp48 is a target of nuclear factor-kappaB activity in thyroid cancer. The Journal of clinical endocrinology and metabolism 2007, 92 (4), 1458–66.

12. Kitange, G. J.; Mladek, A. C.; Schroeder, M. A., et al. Retinoblastoma Binding Protein 4 Modulates Temozolomide Sensitivity in Glioblastoma by Regulating DNA Repair Proteins. Cell reports 2016, 14 (11), 2587–98.

13. Liu, Z.; Li, F.; Zhang, B., et al. Structural basis of plant homeodomain finger 6 (PHF6) recognition by the retinoblastoma binding protein 4 (RBBP4) component of the nucleosome remodeling and deacetylase (NuRD) complex. The Journal of biological chemistry 2015, 290 (10), 6630–8.

14. Lejon, S.; Thong, S. Y.; Murthy, A., et al. Insights into association of the NuRD complex with FOG-1 from the crystal structure of an RbAp48.FOG-1 complex. The Journal of biological chemistry 2011, 286 (2), 1196–203.

15. Schmitges, F. W.; Prusty, A. B.; Faty, M., et al. Histone methylation by PRC2 is inhibited by active chromatin marks. Molecular cell 2011, 42 (3), 330–41.

16. Alqarni, S. S.; Murthy, A.; Zhang, W., et al. Insight into the architecture of the NuRD complex: structure of the RbAp48-MTA1 subcomplex. The Journal of biological chemistry 2014, 289 (32), 21844–55.

17. Murzina, N. V.; Pei, X. Y.; Zhang, W., et al. Structural basis for the recognition of histone H4 by the histone-chaperone RbAp46. Structure 2008, 16 (7), 1077–85.

18. Millard, C. J.; Varma, N.; Saleh, A., et al. The structure of the core NuRD repression complex provides insights into its interaction with chromatin. Elife 2016, 5, e13941.

19. Kuzmichev, A.; Zhang, Y.; Erdjument-Bromage, H., et al. Role of the S¡n3-H¡stone Deacetylase Complex in Growth Regulation by the Candidate Tumor Suppressor p33ING1. Molecular and Cellular Biology 2002, 22 (3), 835–848.

20. Comet, I.; Riising, E. M.; Leblanc, B., et al. Maintaining cell identity: PRC2-mediated regulation of transcription and cancer. Nature reviews. Cancer 2016, 16 (12), 803–810.

21. Ciferri, C.; Lander, G. C.; Maiolica, A., et al. Molecular architecture of human polycomb repressive complex 2. Elife 2012,1, e00005.

22. Schmidberger, J. W.; Sharifi Tabar, M.; Torrado, M., et al. The MTA1 subunit of the nucleosome remodeling and deacetylase complex can recruit two copies of RBBP4/7. Protein science: a publication of the Protein Society 2016, 25 (8), 1472–82.

23. Torchy M. P.; Hamiche A.; Klaholz B. P. Structure and function insights into the NuRD chromatin remodeling complex. Cellular and molecular life sciences: CMLS 2015, 72 (13), 2491-507.

24. Kolla, V.; Naraparaju, K.; Zhuang, T., et al. The tumour suppressor CHD5 forms a NuRD-type chromatin remodelling complex. The Biochemical journal 2015, 468 (2), 345–52.

25. Yao Y. L.; Yang W. M. The metastasis-associated proteins 1 and 2 form distinct protein complexes with histone deacetylase activity. The Journal of biological chemistry 2003, 278 (43), 42560–8.

26. Zhang, Y.; Ng, H. H.; Erdjument-Bromage, H., et al. Analysis of the NuRD subunits reveals a histone deacetylase core complex and a connection with DNA methylation. Genes Dev. 1999, 13 (15), 1924–35.

27. Bowen, N. J.; Fujita, N.; Kajita, M., et al. Mi-2/NuRD: multiple complexes for many purposes. Biochimica et biophysica acta 2004, 1677 (1–3), 52–7.

28. Fujita, N.; Jaye, D. L.; Geigerman, C., et al. MTA3 and the Mi-2/NuRD complex regulate cell fate during B lymphocyte differentiation. Cell 2004, 119 (1), 75–86.

29. Millard C. J.; Fairall L.; Schwabe J. W. Towards an understanding of the structure and function of MTA1. Cancer metastasis reviews 2014, 33 (4), 857–67.

30. Sen N.; Gui B.; Kumar R. Role of MTA1 in cancer progression and metastasis. Cancer metastasis reviews 2014, 33 (4), 879–89.

31. Liu, J.; Wang, H.; Huang, C., et al. Subcellular localization of MTA proteins in normal and cancer cells. Cancer metastasis reviews 2014, 33 (4), 843–56.

32. Kumar R.; Wang R. A. Structure, expression and functions of MTA genes. Gene 2016, 582 (2), 112–121.

33. Manavathi B.; Kumar R. Metastasis tumor antigens, an emerging family of multifaceted master coregulators. The Journal of biological chemistry 2007, 282 (3), 1529–33.

34. Sen N.; Gui B.; Kumar R. Physiological functions of MTA family of proteins. Cancer metastasis reviews 2014, 33 (4), 869–77.

35. Liu, J.; Xu, D.; Wang, H., et al. The subcellular distribution and function of MTA1 in cancer differentiation. Oncotarget 2014, 5 (13), 5153–64.

36. Margueron R.; Reinberg D. The Polycomb complex PRC2 and its mark in life. Nature 2011, 469 (7330), 343–9.

37. Aranda S.; Mas G.;Aranda S. Di CroceL.. Regulation of gene transcription by Polycomb proteins. Science advances 2015, 1 (11), e1500737.

38. Smits, A. H.; Jansen, P. W.; Poser, I., et al. Stoichiometry of chromatin-associated protein complexes revealed by label-free quantitative mass spectrometry-based proteomics. Nucleic acids research 2013, 41 (1), e28.

39. Cao R.; Zhang Y. SUZ12 is required for both the histone methyltransferase activity and the silencing function of the EED-EZH2 complex. Molecular cell 2004, 15 (1), 57–67.

40. Liu, L.; Xu, Z.; Zhong, L., et al. Enhancer of zeste homolog 2 (EZH2) promotes tumour cell migration and invasion via epigenetic repression of E-cadherin in renal cell carcinoma. BJU international 2016, 117 (2), 351–62.

41. Zhang, H.; Qi, J.; Reyes, J. M., et al. Oncogenic Deregulation of EZH2 as an Opportunity for Targeted Therapy in Lung Cancer. Cancer discovery 2016, 6 (9), 1006–21.

42. Hung, S. Y.; Lin, H. H.; Yeh, K. T., et al. Histone-modifying genes as biomarkers in hepatocellular carcinoma. Int J Clin Exp Pathol 2014, 7 (5), 2496–507.

43. Ougolkov A. V.; Bilim V. N.; Billadeau D. D. Regulation of pancreatic tumor cell proliferation and chemoresistance by the histone methyltransferase enhancer of zeste homologue 2. Clinical cancer research: an official journal of the American Association for Cancer Research 2008, 14 (21), 6790–6.

44. Pasini, D.; Bracken, A. P.; Jensen, M. R., et al. Suz12 is essential for mouse development and for EZH2 histone methyltransferase activity. EMBO J 2004, 23 (20), 4061–71.

45. O’Carroll, D.; Erhardt, S.; Pagani, M., et al. The polycomb-group gene Ezh2 is required for early mouse development. Mol Cell Biol 2001, 21 (13), 4330–6.

46. Faust, C.; Schumacher, A.; Holdener, B., et al. The eed mutation disrupts anterior mesoderm production in mice. Development 1995, 121 (2), 273–85.

47. Aloia L.; Di Stefano, B., Di Croce, L., Polycomb complexes in stem cells and embryonic development. Development 2013, 140 (12), 2525–34.

48. Aldiri I.; Vetter M. L. PRC2 during vertebrate organogenesis: a complex in transition. Developmental biology 2012, 367 (2), 91–9.

49. Lee, S. C.; Miller, S.; Hyland, C., et al. Polycomb repressive complex 2 component Suz12 is required for hematopoietic stem cell function and lymphopoiesis. Blood 2015, 126 (2), 167–75.

50. Sun, A.; Li, F.; Liu, Z., et al. Structural and biochemical insights into human zinc finger protein AEBP2 reveals interactions with RBBP4. Protein &, cell 2017.

51. Chen, S.; Jiao, L.; Shubbar, M., et al. Unique Structural Platforms of Suz12 Dictate Distinct Classes of PRC2 for Chromatin Binding. Molecular cell 2018, 69 (5), 840–852 e5.

52. Moody, R. R.; Lo, M. C.; Meagher, J. L., et al. Probing the interaction between the histone methyltransferase/deacetylase subunit RBBP4/7 and the transcription factor BCL11A in epigenetic complexes. The Journal of biological chemistry 2018, 293 (6), >2125–2136.

53. Todd M. A.; Picketts D. J. PHF6 interacts with the nucleosome remodeling and deacetylation (NuRD) complex. Journal of proteome research 2012, 11 (8), 4326–37.

54. Liu, Z.; Li, F.; Ruan, K., et al. Structural and functional insights into the human Borjeson-Forssman-Lehmann syndrome-associated protein PHF6. The Journal of biological chemistry 2014, 289 (14), 10069–83.

55. Lower, K. M.; Turner, G.; Kerr, B. A., et al. Mutations in PHF6 are associated with Borjeson-Forssman-Lehmann syndrome. Nature genetics 2002, 32 (4), 661–5.

56. Bao, Y.; Liu, Z.; Zhang, J., et al. 1H, 13C and 15N resonance assignments and secondary structure of the human PHF6-ePHD1 domain. Biomolecular NMR assignments 2016, 10 (1), 1–4.

57. Voss, A. K.; Gamble, R.; Collin, C., et al. Protein and gene expression analysis of Phf6, the gene mutated in the Borjeson-Forssman-Lehmann Syndrome of intellectual disability and obesity. Gene expression patterns: GEP 2007, 7 (8), 858–71.

58. Zhang, C.; Mejia, L. A.; Huang, J., et al. The X-linked intellectual disability protein PHF6 associates with the PAF1 complex and regulates neuronal migration in the mammalian brain. Neuron 2013, 78 (6), 986–93.

59. Wang, J.; Leung, J. W.; Gong, Z., et al. PHF6 regulates cell cycle progression by suppressing ribosomal RNA synthesis. The Journal of biological chemistry 2013, 288 (5), 3174–83.

60. de Rooij, J. D.; van den Heuvel-Eibrink, M. M., van de Rijdt,N. K., et al. PHF6 mutations in paediatric acute myeloid leukaemia. British journal of haematology 2016, 175 (5), 967–971.

61. Mori, T.; Nagata, Y.; Makishima, H., et al. Somatic PHF6 mutations in 1760 cases with various myeloid neoplasms. Leukemia 2016, 30 (11), 2270–2273.

62. Li, M.; Xiao, L.; Xu, J., et al. Co-existence of PHF6 and NOTCH1 mutations in adult T-cell acute lymphoblastic leukemia. Oncology letters 2016, 12 (1), 16–22.

63. Di Donato N.; Isidor, B., Lopez Cazaux; S., et al. Distinct phenotype of PHF6 deletions in females. European journal of medical genetics 2014, 57 (2–3), 85–9.

64. Zweier, C.; Kraus, C.; Brueton, L., et al. A new face of Borjeson-Forssman-Lehmann syndrome? De novo mutations in PHF6 in seven females with a distinct phenotype. Journal of medical genetics 2013, 50 (12), 838–47.

65. Berland, S.; Alme, K.; Brendehaug, A., et al. PHF6 Deletions May Cause Borjeson-Forssman-Lehmann Syndrome in Females. Molecular syndromology 2011, 1 (6), 294–300.

66. Crawford, J.; Lower, K. M.; Hennekam, R. C., et al. Mutation screening in Borjeson-Forssman-Lehmann syndrome: identification of a novel de novo PHF6 mutation in a female patient. Journal of medical genetics 2006, 43(3), 238–43.

67. Wieczorek, D.; Bogershausen, N.; Beleggia, F., et al. A comprehensive molecular study on Coffin-Siris and Nicolaides-Baraitser syndromes identifies a broad molecular and clinical spectrum converging on altered chromatin remodeling. Human molecular genetics 2013, 22 (25), 5121–35.

68. Jahani-Asl, A.; Cheng, C.; Zhang, C., et al. Pathogenesis of Borjeson-Forssman-Lehmann syndrome: Insights from PHF6 function. Neurobiology of disease 2016, 96, 227–235.

69. Kunze, K.; Gamerdinger, U.; Lessig-Owlanj, J., et al. Detection of an activated JAK3 variant and a Xq26.3 microdeletion causing loss of PHF6 and miR-424 expression in myelodysplastic syndromes by combined targeted next generation sequencing and SNP array analysis. Pathology, research and practice 2014 210(6), 369–76.

70. Todd M. A.; Ivanochko D.; Picketts D. J. PHF6 Degrees of Separation: The Multifaceted Roles of a Chromatin Adaptor Protein. Genes 2015, 6 (2), 325–52.

71. Hajjari, M.; Salavaty, A.; Crea, F., et al. The potential role of PHF6 as an oncogene: a genotranscriptomic/proteomic meta-analysis. Tumour biology: the journal of the International Society for Oncodevelopmental Biology and Medicine 2016, 37 (4), 5317–25.

72. Meacham, C. E.; Lawton, L. N.; Soto-Feliciano, Y. M., et al. A genome-scale in vivo loss-of-function screen identifies Phf6 as a lineage-specific regulator of leukemia cell growth. Genes &, development 2015, 29 (5), 483–8.

73. Fox, A. H.; Liew, C.; Holmes, M., et al. Transcriptional cofactors of the FOG family interact with GATA proteins by means of multiple zinc fingers. EMBO J 1999, 18 (10), 2812–22.

74. Tsang, A. P.; Visvader, J. E.; Turner, C. A., et al. FOG, a multitype zinc finger protein, acts as a cofactor for transcription factor GATA-1 in erythroid and megakaryocytic differentiation. Cell. 1997 Jul 11;90(1)¶. 1997, 90 (1), 109–19.

75. Fox, A. H.; Kowalski, K.; King, G. F., et al. Key residues characteristic of GATA N-fingers are recognized by FOG. J Biol Chem. 1998 Dec 11;273(50) 33595–603.

76. Garriga-Canut M.; Orkin S.H., Transforming acidic coiled-coil protein 3 (TACC3) controls friend of GATA-1 (FOG-1) subcellular localization and regulates the association between GATA-1 and FOG-1 during hematopoiesis. The Journal of biological chemistry 2004, 279 (22), 23597–605.

77. Chlon T. M.; Crispino J. D. Combinatorial regulation of tissue specification by GATA and FOG factors. Development 2012, 139 (21), 3905–16.

78. Fujiwara, T.; Sasaki, K.; Saito, K., et al. Forced FOG1 expression in erythroleukemia cells: Induction of erythroid genes and repression of myelo-lymphoid transcription factor PU.1. Biochemical and biophysical research communications 2017, 485 (2), 380–387.

79. Tsang, A. P.; Fujiwara, Y.; Hom, D. B., et al. Failure of megakaryopoiesis and arrested erythropoiesis in mice lacking the GATA-1 transcriptional cofactor FOG. Genes &, development 1998, 12 (8), 1176–88.

80. Nichols, K. E.; Crispino, J. D.; Poncz, M., et al. Familial dyserythropoietic anaemia and thrombocytopenia due to an inherited mutation in GATA1. Nature genetics 2000, 24 (3), 266–70.

81. Miccio, A.; Wang, Y.; Hong, W., et al. NuRD mediates activating and repressive functions of GATA-1 and FOG-1 during blood development. EMBO J 2010, 29 (2), 442–56.

82. Bagu, E. T.; Miah, S.; Dai, C., et al. Repression of Fyn-related kinase in breast cancer cells is associated with promoter site-specific CpG methylation. Oncotarget 2017, 8 (7), 11442–11459.

83. Katz S. G.; Cantor A. B.; Orkin S. H. Interaction between FOG-1 and the Corepressor C-Terminal Binding Protein Is Dispensable for Normal Erythropoiesis In Vivo. Molecular and Cellular Biology 2002, 22 (9), 3121–3128.

84. Hong, W.; Nakazawa, M.; Chen, Y. Y., et al. FOG-1 recruits the NuRD repressor complex to mediate transcriptional repression by GATA-1. EMBO J. 2005, 24 (13), 2367–78.

85. Song J. J.; Garlick J. D.; Kingston R. E. Structural basis of histone H4 recognition by p55. Genes &, development 2008, 22 (10), 1313–8.

86. Nowak, A. J.; Alfieri, C.; Stirnimann, C. U.; et al. Chromatin-modifying complex component Nurf55/p55 associates with histones H3 and H4 and polycomb repressive complex 2 subunit Su(z)12 through partially overlapping binding sites. The Journal of biological chemistry 2011, 286 (26), 23388–96.

87. Zhang, W.; Tyl, M.; Ward, R.; et al. Structural plasticity of histones H3-H4 facilitates their allosteric exchange between RbAp48 and ASF1. Nature structural & molecular biology 2013, 20 (1), 29–35.

88. Senisterra, G.; Wu, H.; Allali-Hassani, A.; et al. Small-molecule inhibition of MLL activity by disruption of its interaction with WDR5. The Biochemical journal 2013, 449 (1), 151–9.

89. Allali-Hassani, A.; Wasney, G. A.; Siarheyeva, A.; et al. Fluorescence-based methods for screening writers and readers of histone methyl marks. Journal of biomolecular screening 2012, 17 (1), 71–84.

90. Senisterra, G.; Zhu, H. Y.; Luo, X.; et al. Discovery of Small-Molecule Antagonists of the H3K9me3 Binding to UHRF1 Tandem Tudor Domain. SLAS discovery: advancing life sciences R & D 2018, 2472555218766278.

